# Single-cell multiomics of pediatric BM reveals age-dependent differences in lineage differentiation linked to stromal cell heterogeneity

**DOI:** 10.1101/2024.10.31.621155

**Authors:** Evelyn S. Hanemaaijer, Konradin F. Müskens, Ireen J. Kal, Brigit M. te Pas, Patrycja Fryzik, Nina Epskamp, Aleksandra Balwierz, Tito Candelli, Wim J. de Jonge, Thanasis Margaritis, Mirjam E. Belderbos

## Abstract

Childhood is critical for hematopoietic development and the onset of hematologic diseases. To explore hematopoietic changes from infancy through adolescence, we generated a multi-modal single-cell atlas capturing mRNA and surface protein expression of 90.710 bone marrow (BM) cells. This includes hematopoietic stem/progenitor cells and mesenchymal stromal cells, from seven pediatric individuals and two young adults. We demonstrate that young pediatric BM is distinct from adolescents/young adults (AYA), shifting from B-lineage dominance in early childhood to myeloid and T-lineage bias in adolescence. We uncover two distinct lymphoid progenitors (LyPs) subsets regulating this shift: CD127-positive LyPs with B-lineage output, most abundant in early childhood, and CD127-negative LyPs with lymphoid and myeloid features, more common in AYAs. Age-related changes in stromal composition and signaling, mediated by IL-7 and TGF-β1, correspond with this lineage shift. This study provides an in-depth resource for understanding healthy hematologic development and potential early-life perturbations underlying pediatric hematologic diseases.

## Introduction

Hematopoiesis orchestrates the lifelong production of all differentiated blood and immune cells throughout development, adulthood and old age. Due to its close connection to virtually any organ in the human body, the hematopoietic system is a major attribute of systemic health^1–3^. Accordingly, blood production by hematopoietic stem and progenitor cells (HSPCs) is tightly regulated by both cell-intrinsic and extrinsic mechanisms. These mechanisms ensure a balance between HSPC self-renewal and multilineage differentiation to maintain hematopoietic homeostasis, allow rapid responses to stress, and prevent hematopoietic disease.

The composition and function of the hematopoietic system undergo significant changes throughout human life. Extensive research has elucidated key features of human hematopoietic ageing, including a shift from lymphoid-biased to myeloid-biased output^4,5^, an increase in the relative frequency of HSPCs^4^, and a concomitant decline in their regenerative capacity^6–8^. While the differences between young adults and the elderly have been studied in detail^6,9^, there is a relative paucity of studies defining the cellular and molecular composition of hematopoiesis during human childhood^10–12^. Comprehending pediatric hematopoiesis is crucial to understand the development of the human hematopoietic system, as well as pediatric hematologic diseases.

One of the major challenges in characterizing the single-cell composition of pediatric hematopoiesis is obtaining bone marrow (BM) samples from healthy children. Additionally, several hematopoietic and non-hematopoietic cell types critical for hematopoiesis, including HSPCs and mesenchymal stromal cells (MSCs), are rare in bone marrow aspirates, necessitating specific enrichment strategies to capture sufficient cells for in-depth profiling^13^. Consequently, much of our current understanding of pediatric hematopoiesis is based on studies that may not fully capture all relevant cell types. Recent methods to simultaneously measure mRNA and surface protein expression in single cells provide unprecedented opportunities to dissect the composition of hematopoiesis, during steady-state conditions and in hematologic diseases^14,15^. Compared to traditional flow cytometry-based approaches or unimodal single-cell RNA sequencing (scRNA-seq), these methods allow improved cell type identification and more accurate detection of cell states^16^. Furthermore, as mRNA levels do not always correlate with protein expression^16^, the surface protein modality allows validation that specific transcripts of interest are indeed expressed and may have functional consequences.

Here, we performed multi-modal profiling of the transcriptome and cell surface marker expression of single cells derived from the BM of healthy children and young adults. Utilizing scRNA-seq, we comprehensively characterized the transcriptomic profiles of both hematopoietic and non-hematopoietic cell types, identifying age-related differences in cell abundance, cell states and cell-cell communication. In parallel, we employed co-detection of a panel of 138 cell surface proteins to improve dimensional reduction and clustering, substantiate cell type annotation and validate the age-dependent expression of inferred ligand-receptor pairs. We illustrate the utility of our pediatric BM atlas by uncovering various age-related changes in BM cell frequencies, cell states and differentiation trajectories from infancy to adulthood. Importantly, we demonstrate how phenotypic and transcriptional differences within the lymphoid progenitor (LyP) cell population result in B-lineage-biased hematopoiesis in young children. Finally, using interaction analysis, we uncover age-dependent signaling from MSCs to LyPs that may differentially prime them towards stable or B-lineage-biased output. Collectively, this work represents a comprehensive, multimodal pediatric BM atlas that will serve as a valuable reference for future studies on human hematopoietic development and on hematologic diseases, many of which may originate in childhood.

## Results

### Multi-modal single-cell analysis of human bone marrow

To uncover the cellular and transcriptional landscape of human pediatric BM, we established an experimental pipeline that allows for single-cell analysis of rare BM cell types, while maintaining information on cell type frequencies in the original sample. We isolated cells from seven pediatric and two young adult donors, from each of the following three populations: CD235a-(non-enriched, depleted for erythrocytes), CD235a-CD34+ (HSPC-enriched) and CD235a-CD45^low^CD90+ or CD235aCD45^low^CD271+ (MSC-enriched). All individuals served as healthy BM donors for allogeneic hematopoietic cell transplantation (Table 1). For each BM donor, the enriched cell fractions were pooled with the non-enriched cell fraction of a different donor, followed by sequencing of transcriptome and surface protein expression from the same cell, using a customized panel of 138 oligonucleotide-conjugated antibodies and the 10x Genomics platform (Figure 1A). During subsequent data analysis, we assigned each cell to its original sample based on single nucleotide variants (SNVs) unique for each donor. The non-enriched fraction was used to assess the frequencies of cell types present in the BM, while the enriched fractions were used for in-depth assessment of both HSPCs and BM stromal cells.

**Figure 1:**
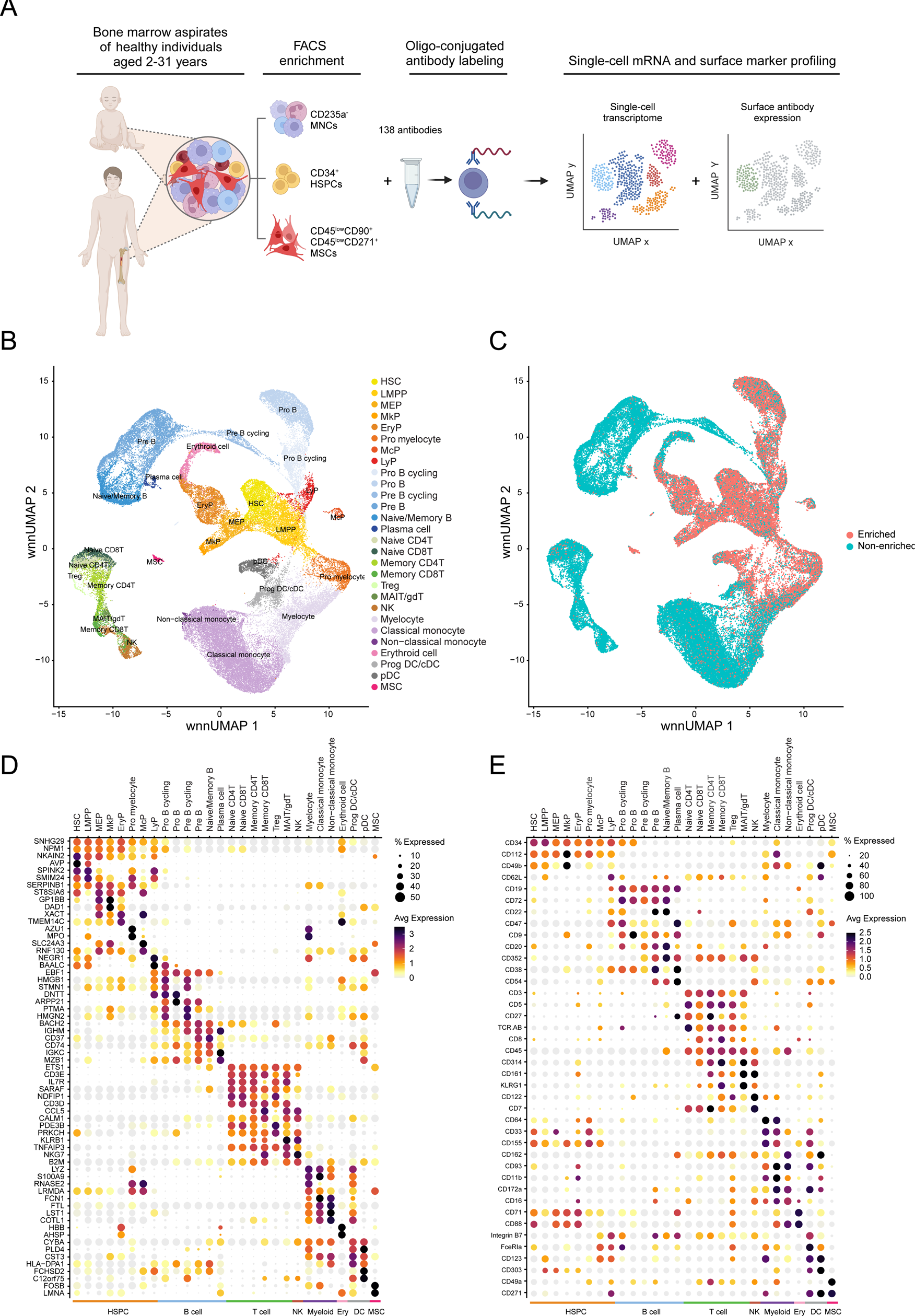
Multimodal single-cell reference atlas of pediatric BM (<18 years). **A)** Overview of the experimental pipeline and cell enrichment strategy. **B**) Weighted nearest neighbor Uniform manifold approximation and projection (wnnUMAP) of single-cell proteo-genomics data of pediatric bone marrow. **C)** UMAP depicting the relative contribution of enriched and non-enriched cell fractions to the dataset. **D-E)** Dot plots showing the top differentially expressed genes (D) and surface markers (E) for each major group and high-resolution cluster. Abbreviations: FACS: fluorescence-activated cell sorting; MNC: mononuclear cell; MSC: mesenchymal stromal cell; HSC: hematopoietic stem cell; LMPP: lympho-myeloid primed progenitor; MEP: megakaryocyte-erythroid progenitor; MkP: megakaryocyte progenitor; EryP: erythroid progenitor; McP: mast cell progenitor; LyP: lymphoid progenitor; Treg: regulatory T cell; MAIT: mucosal-associated invariant T cell; gdT: gmma-delta T cell; NK: natural killer cell; prog DC: dendritic cell progenitor; cDC: conventional DC; pDC: plasmacytoid DC, wnn: weighted nearest neighbor; ADT: antibody-derived tag.

**Table 1:**
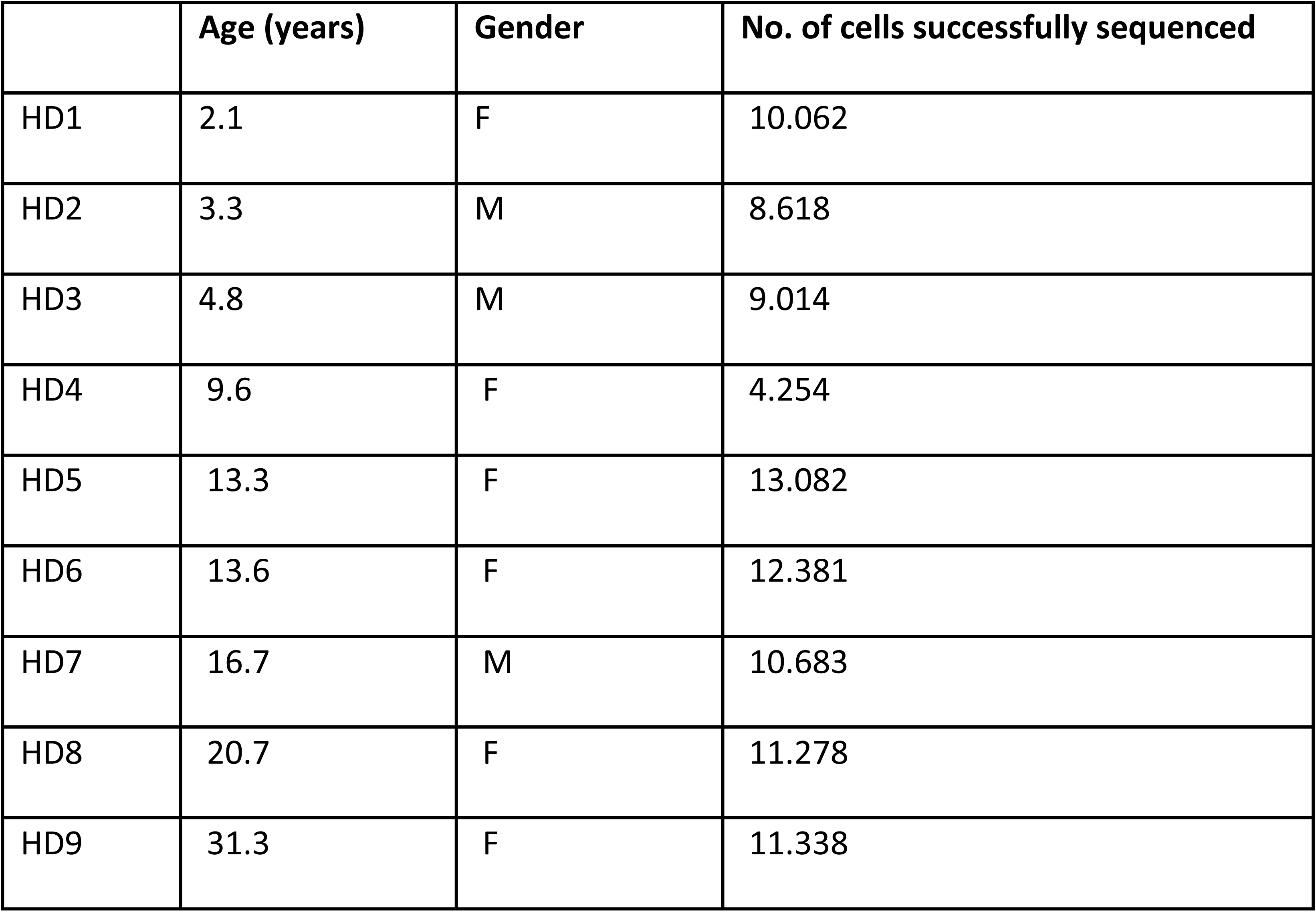
Metadata of donors and samples.

### A single-cell reference atlas of pediatric BM

After quality control, our pediatric dataset comprised of 68.094 cells, with a median of 5133 transcripts and 2001 unique genes per cell for the RNA modality and 1307 counts and 114 unique proteins per cell for the protein modality, respectively. Integration across individuals and measurement modalities revealed 28 distinct cellular clusters (Figure 1B), with each donor contributing to all clusters. Based on comparisons with previously published datasets^10–12^, analysis of marker genes and exploration of the most differentially expressed transcripts (Figure 1D) and proteins (Figure 1E), we annotated these high-resolution clusters and grouped them into eight major groups, representing all major hematopoietic cell lineages. Notably, our enrichment strategy successfully captured large numbers of HSPCs (15.914 cells, 4.9-fold enrichment) and MSCs (570 cells, 18.7-fold enrichment), enabling in-depth investigations (Figure 1C).

### Age-related differences in cellular and transcriptional BM composition

Next, we applied this comprehensive atlas of pediatric BM to explore the developmental changes in BM cell frequencies and transcriptional states from infancy throughout childhood into adolescence. Using the non-enriched cell fractions, we detected a high abundance of B-lineage cells in BM of individuals aged nine years and younger, occupying the majority of total BM cellularity (Figure 2A). In contrast, the BM composition of donors aged 13 years and older had more variation between donors and was predominantly composed of myeloid and T cells. To validate whether the observed differences persisted into young adulthood, we expanded our dataset to include two adult BM donors (20.7 and 31.3 years old; 11.278 and 11.338 cells respectively, Table 1). BM composition in these adult individuals closely resembled that of the adolescent donors (Figure 2A), which was further supported by unsupervised analysis of BM composition across individuals (Figure 2B). For further analysis, BM samples were grouped into two age categories: young pediatric individuals (aged 2-9 years), and adolescents/young adults (AYAs, aged 13-31 years, Figure 2B). In general, young pediatric individuals had significantly higher frequencies of B-lineage cells (median 49.7% vs 13.9%, P=0.016). In contrast, AYA BM had a high abundance T-lineage cells (median 18.6% vs 10.4%, P=0.56) and myeloid lineage cells (median 51,7% vs 24,0%, P=0.063) (Figure 2C). Overall, these data are in line with previous observations^17^. These findings demonstrate that human young pediatric BM is distinct from that of adolescents and young adults and reveal a systemic shift from B-lineage bias towards increased T-cell and myeloid output from human infancy through young adolescence.

**Figure 2:**
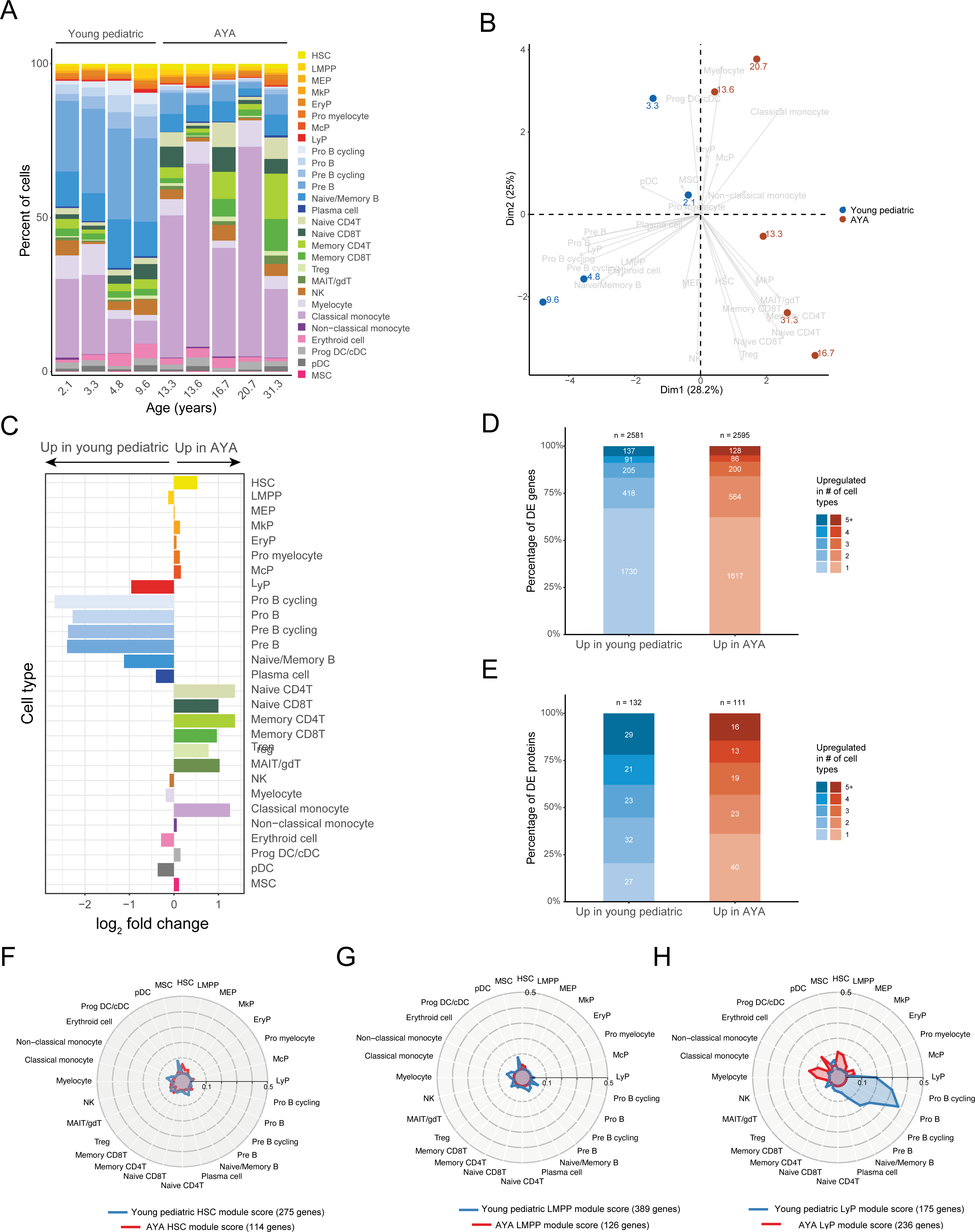
Age-dependent differences in BM composition and gene expression. **A)** Stacked bar plots depicting the cellular composition of the non-enriched cell fraction of seven pediatric (left) and two young adult (right) bone marrow aspirates. **B**) BiPlot visualizing Principal component (PC) analysis of BM composition. Colored dots depict PC scores of individual samples. Grey arrows depict vectors reflecting the contribution of each cluster’s abundance to the principal components. **C)** Bar plot showing the relative abundance of BM cell populations in adolescent and young adult (AYA) versus young pediatric BM. Negative values indicate increased cell type frequencies in young pediatric BM. **D-E)** Stacked bar plots showing the number of differentially expressed genes (D) and surface markers (E) between young pediatric and AYA cell types, categorized by the number of cell types in which they are identified. **F-H)** Radar plots showing the average expression of age-specific modules across all cell types. For each cell type, HSCs (F), LMPPs (G) and LyPs (H), genes upregulated in young pediatric (blue) or AYA (red) cells were grouped into a single module score, and average expression of this module score was measured across all cell types.Abbreviations: HSC: hematopoietic stem cell; LMPP: lympho-myeloid primed progenitor; MEP: megakaryocyte-erythroid progenitor; MkP: megakaryocyte progenitor; EryP: erythroid progenitor; McP: mast cell progenitor; LyP: lymphoid progenitor; Treg: regulatory T cell; MAIT: mucosal-associated invariant T cell; gdT: gamma-delta T cell; NK: natural killer cell; prog DC: dendritic cell progenitor; cDC: conventional DC; pDC: plasmacytoid DC, MSC: mesenchymal stromal cell, log2FC: log-2 fold change.

To explore whether the observed age-related differences in cell frequencies corresponded to functional changes in these populations or their upstream progenitors, we compared single-cell transcriptomes and surface marker expression between young pediatric and AYA BM. Specifically, we aimed to determine whether these shifts in cell type composition reflected a general ageing program active across all cell types, or if age-related changes were more specific to certain cell types or lineages. For each cluster, we performed differential expression between cells from the young pediatric and AYA groups, identifying large numbers of differentially expressed genes (Figure 2D) and proteins (Figure 2E). The majority of age-specific, differentially expressed genes were unique to individual cell types (67% of genes upregulated in pediatric BM and 62% of genes upregulated in AYA BM, respectively, Figure 2D-E). However, certain age-related genes were shared across multiple cell types, potentially reflecting the presence of overarching age-related transcriptional programs. Among these shared genes were those previously linked to ageing in adults (e.g., CTSW across cell lineages^18^, KLF6 in HSPCs^19^), suggesting that certain ageing-related transcriptional programs may initiate in childhood. Notably, the cell-type specificity of most age-related changes, which may not be captured in bulk samples, emphasizes the importance of studying hematopoietic ageing mechanisms in sorted cell populations or at the single-cell level.

To assess whether the transcriptomic differences between young pediatric and AYA BM cells were related to lineage bias, we calculated module scores for each cell type based on the differentially expressed genes between these two groups. We then assigned these scores to each cell in our dataset. Subsequently, we plotted the average expression of these modules across all cell types, focusing on the trajectory from HSCs towards the B-cell lineage (Figure 2F-H). The gene modules of HSCs and LMPPs were equally expressed across all cell types, indicating that these sets of age-dependent differentially expressed genes are not associated with lineage bias. In contrast, the module score for genes upregulated in young pediatric LyPs was strongly enriched in B-lineage cells, while module score upregulated in AYA LyPs were enriched in early progenitors and myeloid cells, indicating that differential priming towards B-versus balanced/myeloid output is evident at the LyP level. Together, these analyses reveal significant developmental changes in the cellular and transcriptional landscape of human BM from childhood into adolescence/young adulthood, with a general B-lineage bias in young pediatric BM, which might be regulated at the level of LyPs.

### Age-related heterogeneity in lymphoid progenitor subsets

Given the higher abundance of LyPs in young pediatric BM and the observed age-related differentiation bias in LyPs, we sought to further dissect the heterogeneity within this cell population. We subclustered the LyP, revealing two distinct populations with distinct transcriptomic and proteomic features (Figure 3A). The first cluster consisted of cells expressing high levels of genes involved in B-lineage differentiation such as EBF1, CD79A and VPREB1, and high surface expression of CD127 (IL-7 receptor, Figure 3B and 3C). We therefore labelled this cluster LyP B-biased (LyP-B). The second cluster consisted of cells expressing high levels of genes associated with stemness (e.g., SPINK2, ABCB1, LRMDA), myeloid (LGALS1) and lymphocyte lineage markers (CD37). Cells in this cluster also displayed increased surface protein expression of markers associated with myeloid lineage (e.g., CD123) and lymphoid lineage (e.g., CD18). We therefore labelled this cluster LyP stable (LyP-S). The differential lineage bias between these LyP subsets was reflected by Gene Ontology term enrichment analysis, with LyP-Bs expressing pathways related to B cell receptor signaling, lymphocyte proliferation and differentiation, and LyP-S expressing pathways related to B-cell activation, as well as myeloid leukocyte activation and mononuclear cell differentiation (Figure 3D). Finally, inference of gene regulatory networks demonstrated increased activity of B lineage-associated regulons in LyP-B (e.g., PAX5, LEF1 and TCF3)^20,21^, while the most differentially active regulons in LyP-S were related to general lymphoid development and stemness (e.g., ELF4, RXRA, Figure 3E)^22,23^.

**Figure 3:**
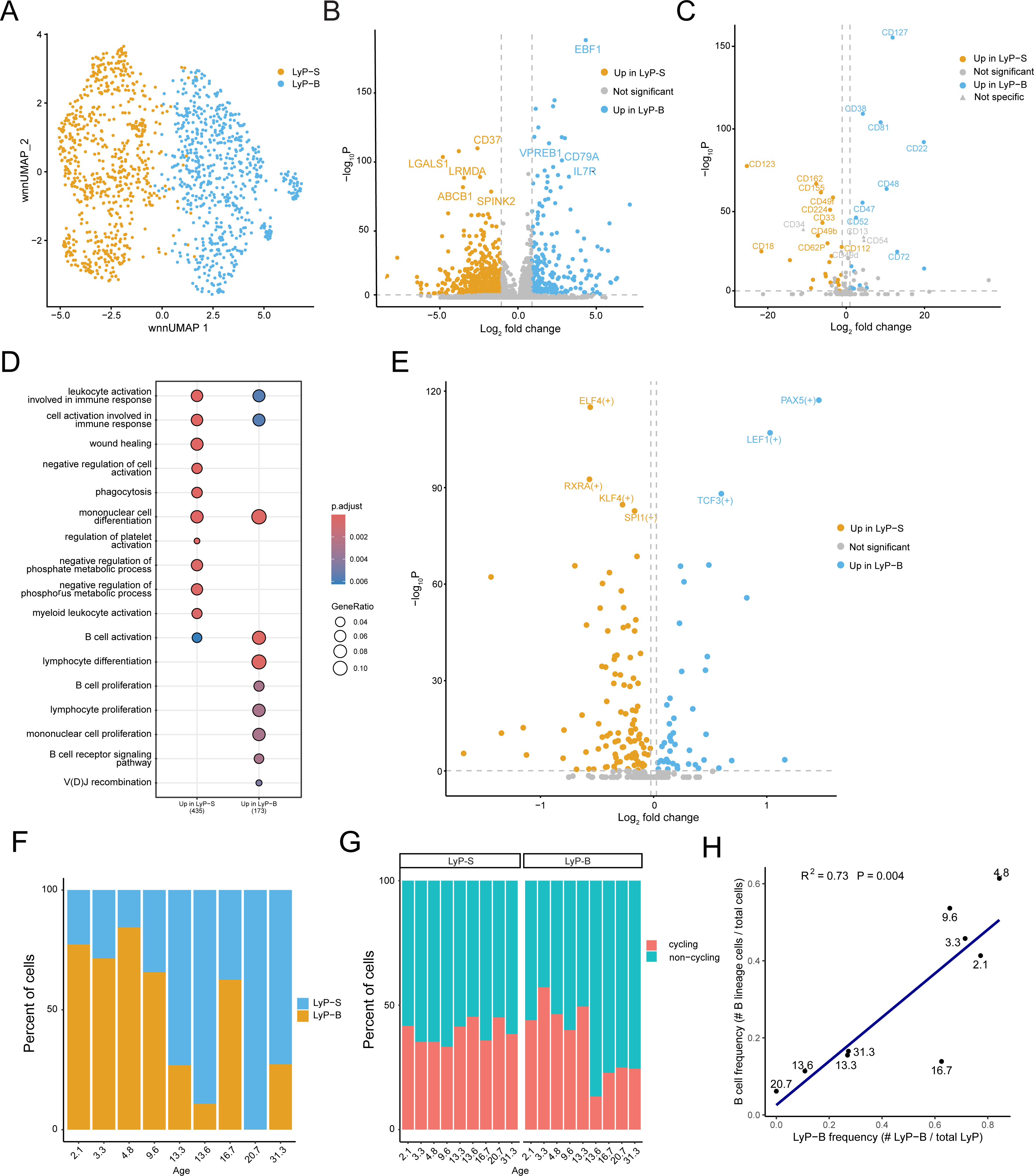
In-depth analysis of lymphoid progenitor subsets. **A)** UMAP lymphoid progenitors (LyPs, n=1339. **B)** Volcano plot depicting differentially expressed genes between B-biased LyP (LyP-B, blue) and stable LyP (LyP-S, orange). **C)** Volcano plot showing differentially expressed surface markers between LyP-B and LyP-S. **D)** Gene ontology enrichment analysis of genes upregulated in LyP-B and LyP-S. **E)** Volcano plot depicted differentially active regulons, calculated using Single cell rEgulatory Network Inference and Clustering in python (pySCENIC). **F)** Stacked bar plots showing relative proportions of LyP-B and LyP-S subsets within total LyP population (non-enriched) for each individual. **G)** Stacked bar plots showing cell cycle assignment of LyP-S (left) and LyP-S (right) subsets (non-enriched + enriched) per individual. **H)** Correlation plot, depicting the correlation between the relative abundance of LyP-B versus the percentage of B-lineage cells (non-enriched) in BM. Each dot represents one individual.

Next, we set out to explore whether differences in the relative abundance of these distinct LyP subsets could contribute to the observed B-lineage bias in young children compared to AYAs. Notably, while all individuals contained both LyP subsets, LyP-B were significantly more abundant in young pediatric BM, while LyP-S were more abundant in AYA BM (median 74.4% vs 26.9% LyP-B, P=0.016, Figure 3F). Of note, in some adult individuals, LyP-B could only be detected in the enriched cell fraction, supporting the relevance of our enrichment approach. Cell-cycle analysis demonstrated that the percentage of cycling cells (S + G2/M) was high in pediatric LyP-B versus LyP-B of AYAs (median 45.2% vs 24.5%, P=0.11, Figure 3F), which may contribute to their increased abundance in young pediatric BM (Figure 3G). Moreover, the frequency of LyP-B in each individual was significantly correlated with the overall percentage of B-lineage cells (R^2^ 0.73, p<0.004) (Figure 3H). Thus, in line with previous reports^17,24^, we provide evidence for the existence of two phenotypically and functionally distinct LyP subpopulations. In addition, we demonstrate here that the abundance of these subsets is age-dependent and associated with a systemic bias towards B-lineage cells in young pediatric BM.

### Age-dependent LyP-bias correlates to developmental heterogeneity in BM stromal cells

Next, we asked whether age-dependent interactions between LyP subsets and their surrounding BM cell types could explain the observed difference in the abundance of LyP-B versus LyP-S between pediatric and AYA donors by affecting LyP specification. To this end, we used NicheNet to predict intercellular communication, linking differentially expressed target genes to potential regulatory ligands^25^. We prioritized expressed ligands based on the predicted regulatory influence on the differentially expressed genes between LyP-B and LyP-S, and the differential expression of their cognate receptor between LyP-B and LyP-S. We first focused on genes upregulated in LyP-B (Figure 3B, Figure 4A, top panel), and identified IGF2, IL-7 and BTLA as the top candidate ligands driving the transcriptional changes in LyP-B. Among these, IL-7 appeared the most promising, showing strong regulatory potential on key markers of LyP-B fate, including EBF1, LEF1, and PAX5. For genes upregulated in LyP-S, we identified TGF-β1 as potential regulator of LyP-S specification (Figure 4A, bottom panel). Interestingly, IL-7 also showed regulatory potential of LyP-S target genes, while TGF-β1 also exhibited strong regulatory potential of LyP-B target genes, suggesting that these ligands may promote one LyP subset while suppressing the other.

**Figure 4:**
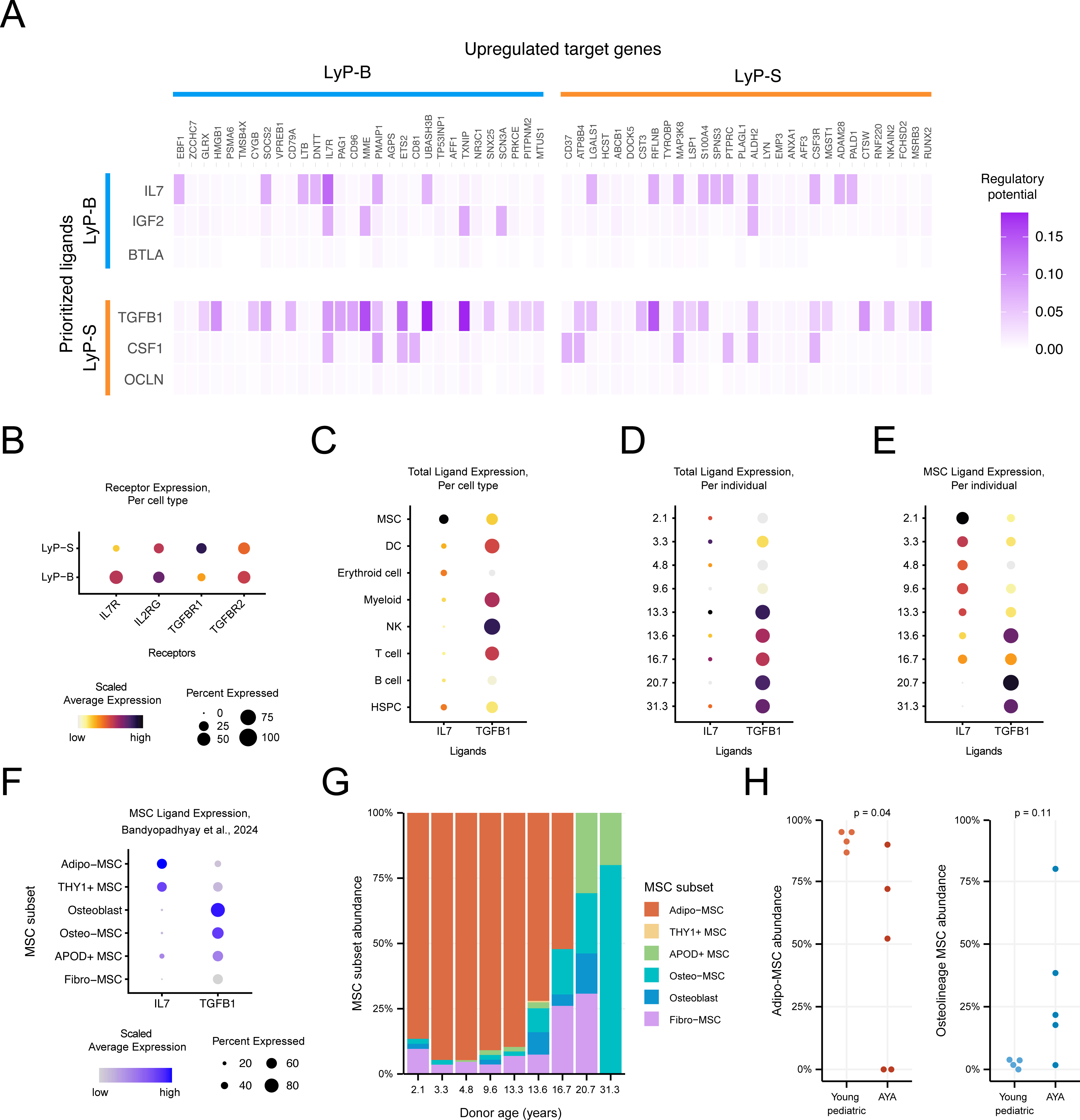
Analysis of putative interactions underlying LyP subset specification. **A)** NicheNet interaction analysis, depicting the regulatory potential of the top 3 predicted ligands underlying the observed upregulation of either LyP-B genes (top rows) and LyP-S genes (bottom rows), prioritized based on regulatory potential and differential receptor expression between LyP subsets. Genes shown are the top 30 differentially expressed genes in LyP-B (left) or LyP-S (right) that are regulated by these ligands. Regulatory potential of LyP-B ligands on LyP-S genes (top right quadrant) and vice versa (bottom left quadrant) are also shown. **B)** Dot plot showing the average expression of receptors for IL-7 signaling and TGF-β1 signaling in LyP-B and LyP-S subsets, scaled across all cell types. **C-E)** Dot plots showing the average expression of IL-7 and TGF-β1 either per cell type (C) or per individual in the total sample (D) or in MSCs only (E). Expression is scaled across all cell types (C), all samples (D) or all MSCs (E). **F)** Dot plot showing the expression of IL-7 and TGF-β1 in different MSC subsets in an external MSC dataset^26^, scaled across all MSC subsets. **G)** Bar plots showing the relative contribution of different MSC subsets to the total MSC population in each individual. **H)** Comparison of the relative contribution of adipo-MSCs and osteolineage MSCs to the total MSC population in young pediatric and AYA BM aspirates. Abbreviations: LyP: lymphoid progenitor; MSC mesenchymal stromal cell; NK: natural killer cell; DC: dendritic cell; HSPC: hematopoietic stem and progenitor cell.

To further delineate the identified interactions, we quantified receptor expression in both LyP subsets. LyP-B showed significantly increased expression of IL-7R (Fold Change [FC]: 7.01, P<0.001) and its co-receptor IL-2Rγ (FC: 1.45, P= 0.002), while LyP-S showed significantly elevated expression of TGFBR1 (FC: 2.26, P<0.001, Figure 4B). TGFBR2 expression was not different between LyP-B and LyP-S (FC = 1.3, P= 0.45). Since the IL-7R (also known as CD127) was included in our ADT panel, we also confirmed the differential expression of this receptor at protein level (Figure 3C). We next examined the expression of each ligand in our reference map. While TGF-β1 was broadly expressed across multiple cell types, IL-7 expression was mostly restricted to MSCs (Figure 4C). Both ligands showed age-dependent expression patterns, with IL-7 expression peaking in early infancy and gradually decreasing through adolescence, while TGF-β1 expression steadily increased with age, reaching its highest level in AYAs (Figure 4D-E). Therefore, the age-dependent expression of both ligands and receptors matches the observed age-dependent differences in LyP subset abundance, further suggesting a direct regulatory role for IL-7 and TGF-β1 in instructing LyP differentiation.

Recent adult human stromal cell atlases have identified distinct MSC subsets expressing varying levels of HSPC signaling molecules. Therefore, we next explored whether age-dependent differences in IL-7 and TGF-β1 expression were related to age-dependent changes in the abundance of these subtypes. Using the MSC scRNA seq data from one of these atlases^26^, we found that IL-7 expression was highly expressed in adipo- and THY1+ MSCs and low IL-7 in osteoblast and osteo-MSCs, while TGF-β1 was highest in osteolineage MSCs, indicating MSC subtype-specific expression patterns (Figure 4F). We then performed label transfer to annotate the MSCs in our dataset according to these subtypes (Figure 4G). Notably, the proportion of IL-7-expressing adipo-MSCs significantly decreased with age (P=0.04), as the abundance of TGF-β1 expressing osteolineage MSCs (Osteo-MSC and Osteoblast) increased (P=0.11) (Fig. 4G-H). Together, these findings identify previously unrecognized age-dependent changes in BM stromal composition that may contribute to the altered LyP-B ratios observed between young pediatric and older individuals.

## Discussion

Here, we present a comprehensive single-cell analysis of pediatric BM, with a particular focus on HSPCs and the stromal niche. We uncover dynamic age-related changes in BM cellular composition, differentiation trajectories, and microenvironmental interactions from infancy through adolescence. Importantly, we demonstrate that the differential bias towards production of B-lineage cells in early childhood is linked to age-related differences in distinct LyP subsets, likely driven by reciprocal interactions with the BM niche. In support of this, we unveiled age-dependent heterogeneity in BM MSCs from infancy through adolescence/young adulthood, with increased frequencies of adipo-MSCs in young children, capable of instructing B-lineage transcriptional programs in LyPs through IL-7/IL-7R signaling.

Our data provide a high-resolution atlas of the single-cell transcriptome and surface marker expression profiles in healthy pediatric BM, providing an important reference for studies on pediatric hematologic disorders. Given the growing importance of single-cell maps in understanding normal and malignant hematopoiesis, our pediatric-specific dataset addresses a critical gap in understanding BM biology across different stages of human development. In addition, this study offers key insights into age-specific regulatory mechanisms that may contribute to the age-dependent occurrence of hematologic malignancies. Here, we highlight some of these insights.

First, we identified two transcriptionally distinct subpopulations of LyPs, which differ in surface marker expression, lineage differentiation programs, and age-dependent prevalence. Previous studies have identified similar heterogeneity in LyP subsets, demonstrating that CD127^-^ and CD127^+^ early LyPs originate from a common multipotent lymphoid progenitor (MLP), but exhibit differential polarization towards natural killer (NK)/innate lymphoid cells (ILC)/T or B-cell lineages. While FLT3 signaling was suggested to primarily drive CD127^-^ differentiation, the emergence and proliferation of CD127^+^ LyPs was previously considered cell-autonomous. Our study confirms and builds upon these findings by identifying niche-derived IL-7 as a cell-extrinsic regulator of CD127^+^ LyPs, influencing the age-dependent bias between NK/ILC/T and B lineages.

Second, this study establishes a framework for understanding transcriptional plasticity of the human HSPC compartment. Surprisingly, we found several myeloid-associated transcription factor programs (e.g., CEBPA, GATA2) to persist into the LyP stage. This retention of myeloid potential within human LyPs is supported by recent independent studies^27,28^, suggesting that lineage restriction at the LyP stage might be less stringent than previously thought. This finding has significant implications for lineage differentiation during normal and stress hematopoiesis, as well as for the potential of B-cell malignancies to adapt their transcriptional program and undergo lineage switching upon selective pressure.

Third, our work demonstrates that the composition of the BM stroma and its instructive signals towards hematopoietic cells undergo substantial changes throughout pediatric development that may be linked to age-dependent differences in hematopoietic function. Previous studies have been instrumental in identifying functionally heterogeneous MSC subsets within the human BM niche and characterizing their transcriptomic and functional features. Building upon these data, we demonstrate that the frequency of distinct MSC subsets is age-dependent, with increased frequencies of IL-7 producing adipo-and THY1+ MSCs in young individuals. Of note, since we used BM aspirates as a sample source, we may not have captured the full spectrum of MSC subpopulations or their spatial context, which should be further investigated.

Finally, the developmental states observed within the healthy pediatric lymphoid lineage closely resemble the recent findings on transcriptomic heterogeneity within B-ALL^27^. Interestingly, bulk leukemic samples from young children displayed a stronger B-lineage transcriptional signature compared to those diagnosed in adolescents/young adults, which had more multipotent transcriptional profiles. Our data demonstrate that these age-dependent transcriptional programs mirror the age-dependent transcriptomic states of the LyP population. Since the cell of origin for pediatric B-ALL is thought to emerge years before clinical diagnosis, perhaps even during fetal development^29–31^, these findings suggest that the capacity of the BM niche to instruct B-lineage differentiation is preserved in the malignant setting. Accordingly, elevated levels of IL-7 in pediatric BM, which has been shown to induce a preleukemic state^31,32^, may facilitate the acquisition of further genetic or molecular aberrations that drive B-cell leukemogenesis. Given the increasing application of lymphoid-targeted therapies and the potential for lineage switch as a mechanism of escape, harnessing lineage-instructive signals from the BM niche to affect B-ALL transcriptome may provide a currently unexplored therapeutic strategy.

In conclusion, our single-cell pediatric BM map provides relevant insights into the ontogeny of the human hematopoietic system and its age-dependent modulation by specific stromal cell populations. By allowing for age-matched comparisons, this resource will facilitate a deeper understanding of hematological diseases originating in childhood and provides a foundation for exploring tumor-stromal interactions as a therapeutic target for pediatric leukemias.

## Methods

### Human samples

Bone marrow (BM) aspirates from healthy individuals were acquired through the Biobank of the Princess Máxima Center for Pediatric Oncology, Utrecht, the Netherlands. BM was collected bilaterally from the posterior superior iliac crests, as part of a stem cell donation to an affected relative. None of the healthy donors received any HSC-mobilizing treatment prior to stem cell donation. All individuals (and if minor, their legal representatives) provided informed consent. After graft infusion, residual BM mononuclear cells from the aspirate were isolated by Ficoll (Cytiva Life Sciences) density gradient centrifugation and cryopreserved in liquid nitrogen until further use.

Ethical approval for this study was obtained from the Institutional Review Board of the Princess Máxima Center (PMCLAB 2022.0328) in accordance with the Declaration of Helsinki. Donor demographic data and sample details are provided in **Table 1**.

### Sample preparation

Cryopreserved bone marrow samples were thawed rapidly in a water bath at 37°C. An equal volume of prewarmed thawing medium (DMEM, high glucose, pyruvate, no glutamine, 20% fetal calf serum (FCS)) was added to the viably frozen cell suspension in a dropwise fashion. The cell suspension was transferred to a 50 ml falcon tube and was further diluted (1:10) by dropwise addition of pre-warmed thawing medium. Cells were centrifuged at 400 *g* for 5 minutes at 4°C. The cell pellet was resuspended in thawing medium containing DNAse (100 µg/ml, Roche) with MgCl_2_ (10 mM, Merck) and incubated for 30 minutes at 4°C. After incubation, cells were centrifuged at 400 *g* for 5 minutes at 4°C and resuspended in cell staining buffer (CSB; Biolegend). The number of cells in the suspension was established using the Countess^TM^ II cell counter (Invitrogen).

### Cell sorting and multiplexing for CITE-Seq

Prior to library preparation, each individual’s sample was enriched for HSPCs and MSCs, which were multiplexed with the non-enriched cell fraction of another genetically distinct individual. To this aim, FcR blocking reagent (Human Trustain FcX, Biolegend) was added to the cells at a 1:10 dilution and incubated on ice for 5 minutes. Cells were then incubated with Zombie NIR^TM^ viability dye (BioLegend), on ice in the dark for 15 minutes. After the incubation, the cell suspension was washed by adding CSB and centrifuging at 400 *g* at 4°C, followed by resuspension. Next, the cells were incubated on ice in the dark for 30 minutes with a customized mix of fluorophore-conjugated antibodies and oligonucleotide-conjugated antibodies. Following incubation, the cells were washed three times with CSB and centrifuged at 400 *g* at 4°C after each wash. The cell suspension was then filtered using a 35 µM filter and sorted using the Sony SH800S cell sorter (SONY SH800S system software v2.1), with a uniform gating strategy for all samples. In general, we sorted 2.5×10^4^ non-enriched, erythrocyte precursor-depleted cells (Zombie NIR^-^, CD235a^-^) from one individual, and combined with 1×10^4^ HSPCs (Zombie NIR^-^CD235a^-^CD45^-^CD34^+^) and up to 2×10^3^ MSCs (Zombie NIR^-^ CD235a^-^CD45^low^CD34^-^CD271^+^ or Zombie NIR^-^CD235a^-^CD45^low^CD34^-^CD90^+^) of another genetically distinct individual. During subsequent data analysis (described below), cells were demultiplexed and assigned to their original sample based on SNVs specific for each individual. After mixing, the resulting cell suspensions were counted using Trypan Blue and a Bürker counting chamber.

### Library preparation and sequencing

Approximately 10.000 cells per multiplexed sample were loaded onto a Chromium Single Cell G chip and used for library preparation using a Chromium Next GEM Automated Single Cell 3’ Library and Gel Bead Kit v3.1 (10x Genomics) according to the manufacturer’s instructions. For each multiplexed sample two library were prepared, one for the RNA and one for the antibody capture modality. Each library was sequenced using the NovaSeq 6000 (Illumina), using the following number of cycles: Read 1: 28, i7: 10, i5: 10 and Read2: 91.

## CITE-Seq data analysis

### Pre-processing

CITE seq data were processed using CellRanger count with feature barcoding (version 7.1.0, 10X Genomics), using the refdata-gex-GRCh38-2020-A transcriptome and a modified Feature Reference file.

### Genotype demultiplexing

Cells from multiplexed samples were SNV-based genotype-demultiplexed using souporcell (Singularity image created 1 December 2021)^33^.

### Barcode filtering

Barcodes with less than 1500 transcripts, and/or a percentage of mitochondrial genes above 10% were removed. Also, barcodes classified as doublets or unassigned genotypes by souporcell^33^, and barcodes classified as doublets in over 5 out of 10 runs using scDblFinder (version 1.18.0)^34^ were discarded.

### Normalization, dimensional reduction, feature deconfounding and integration

Further processing and analyses were carried out using Seurat (version 5.1.0)^35^. For each individual, donor gene expression data were SCTransform-normalized with SCTransform (v2, number of variable features = 3000). Dimensional reduction was conducted with the RunPCA function from Seurat. SCT integration features were calculated using the SelectSCTIntegrationFeatures function from Seurat.

Using gene lists from the Scutils package (version 1.123) the following genes were filtered out from the SCT integration features, provided they were also found as variable features: genes specific to S- or G2M cell cycle phases, donor specific genes correlating with S- or G2M phases, male and female-specific genes, stress-related genes and ribosomal protein genes. Gene expression data was integrated across donors with canonical correlation analysis (CCA), using the IntegrateLayers function in Seurat, using the filtered SCT integration features. Antibody capture data were normalized per library using DSBantibody normalization (version 1.0.3)^36^. For each individual donor, FindVariableFeatures and ScaleData were run and dimensional reduction was done with the RunPCA function from Seurat. Data were also integrated with CCA using the IntegrateLayers function in Seurat, using all antibodies as integration features.

### Visualization, clustering and cell type annotation

A weighted-nearest-neighbor (WNN) graph was created with the FindMultiModalNeighbors function in Seurat, using the integrated reductions of both RNA and ADT modalities. A WNN UMAP was created from this WNN graph with RunUMAP, using 30 principal components for both modalities. Clustering was done using FindClusters from Seurat with the wsnn graph, the SLM modularity optimization algorithm, as recommended by the Weighted Nearest Neighbor Analysis vignette from Seurat, and resolution=0.4.

Cell type annotation was performed by combining three complimentary annotation approaches, performed with gene expression log-normalized data. First, cell types were inferred with SingleR (version 2.6.0)^36^. Second, individual cells were mapped to two external reference BM datasets^11,12^. Mapping to the CITE-seq dataset bmcite (version 0.3.0) from SeuratData (version 0.2.2.9001) was done as recommend by the Seurat multimodal reference mapping vignette^37^. The Gene Expression bone marrow dataset in Deeply Integrated human Single-Cell Omics (DISCO)^38^ was down-sampled to a maximum of 3000 cells per cell type and SCTransform normalized, followed by generation of a PCA and UMAP, which were in turn used for reference mapping, according to the Seurat Mapping and annotating query datasets vignette. Third, antibody capture data and RNA expression data from known marker genes were used to confirm cell type annotations.

For in-depth analysis of the erythroid lineage, the myeloid lineage, and for T and NK cells, the respective clusters were subsetted, followed by subset-specific SCT normalization, dimensional reduction, feature deconfounding, integration and clustering at resolutions of 0.7, 0.5 and 0.3, respectively. Cell type annotations were then redefined as described above. The resulting Seurat object provided cell type annotations at multiple levels of resolution, ranging from broad categories (major groups, e.g., T-cells) to more detailed classifications (high-resolution clusters, e.g. gd T-cells).

### Subclustering of LyP

For in-depth analysis of LyPs, these cells were subsetted and analysed as above, with the following deviations: for clustering a resolution of 0.1 was used; CCA integration between donors was not performed.

### Differential gene and protein expression

To compare each major groups and high-resolution cell type against all other cell types, we used the wilcoxauc function in presto (version 1.0.0) to find cell-type-specific, significantly differentially expressed genes and proteins. We used the following thresholds: p-adj <0.01, pct_in - pct_out >= 0, pct_in > 20, auc > 0.5, logFC > 0. For each cell type, the FindMarkers function (test.use = “wilcox”, logfc.threshold = 0, min.pct = 0.01) was used to identify genes and proteins with significant differential expresseion between young and old cells (thresholds: p-adj <0.01 & FC > 1.5 and pct.1 > pct.2, or p-adj <0.01 & FC < −1.5 and pct.1 < pct.2). From these cell-type-specific young and old gene lists, we computed module scores using Seurat’s AddModuleScore function and assessed scores per cell. Module scores were visualized as mean module scores per cluster using the ggradar package (version 0.2). For LyP subclusters, the FindMarkers function (test.use = “wilcox”, logfc.threshold = 0, min.pct = 0.01) was used to identify significantly differentially expressed genes and proteins between cluster 0 and cluster 1 (thresholds: p-adj <0.01 and FC > 2 & pct.1 > pct.2, or p-adj <0.01 & FC < −2 and pct.1 < pct.2). For all differential gene expression analyses, male and female-specific genes were removed. For comparisons of major groups and high-resolution clusters, confounder genes mentioned above were also removed. Gene Ontology enrichment analysis was done using the enrichGO function from the clusterProfiler (version 4.12.0) package, using Biological Process terms.

### Transcription factor activity analysis

Inference of transcription factor network activity was done using Single-Cell rEgulatory Network Inference and Clustering in python (pySCENIC, image version aertslab-pyscenic-0.11.2.sif)^39^. For this, we used a loom file with raw transcript counts of a downsampled dataset as input, along with the “hs_hgnc_tfs.txt” transcription factor list, the “motifs-v9-nr.hgnc-m0.001-o0.0.tbl” motifs, and the “hg38 refseq-r80 10kb_up_and_down_tss.mc9nr.feather” input databases. Only transcription factors that were identified in ≥2 out of 3 independent runs were analyzed. Activity per cell was calculated as the mean AUCell values across runs. For LyP subclusters, FindMarkers (test.use = “wilcox”, logfc.threshold = 0, min.pct = 0.01) was used to identify significant differential regulon activity between cluster 0 and cluster 1 (thresholds: p-adj <0.01, FC > 1).

### Cell interaction analysis

Interaction analysis and ligand prioritization were performed using NicheNet (nichenetr package, version 2.2.0)^25^. Ligand activity analysis was performed on genes upregulated in LyP-B or genes upregulated in LyP-S (identified using the FindMarkers function as described above), using the top n=5000 downstream targets for each ligand and a quantile cutoff of 0.001. Ligands were prioritized by equally weighing the following two criteria: (1) the predicted ligand activity; and (2) LyP subset-specific receptor expression.

### MSC subtype identification and analysis

For identification of MSC subtypes, the enriched and non-enriched MSCs were pooled, subsetted, mapped and annotated, following the Seurat Mapping and annotating query datasets vignette, using an adult human stromal cell atlas^13^.

### Statistical analyses

Dimensional reduction of BM composition data was performed using principal component analysis, using the prcomp function from the stats package (version 4.4.0). The relative abundance of cell types, LyP-B and LyP-S subsets, cycling vs non-cycling LyPs and MSC subsets between young pediatric BM and AYA BM was compared using Wilcoxon rank sum test. To allow reliable comparisons of rare cell types (cycling vs non-cycling LyPs and MSC subsets) the enriched and non-enriched were pooled. For all other comparisons, the non-enriched fractions were used.

## Acknowledgements

We thank the study subjects and their guardians for facilitating this research. We thank Philip Lijnzaad and Jeff DeMartino for direct technical assistance. We acknowledge the financial support provided by the European Hematology Association (EHA; Physician Scientist Grant), the Dutch Research Council’s Veni Grant (#VI.Veni.202.021), the Landsteiner Foundation for Blood Transfusion Research (LSBR; #2305F), and the European Research Council (ERC; #101114895).

## Data Availability

Requests for access to the data supporting the findings of this study can be addressed to the corresponding author.

